# Application of machine learning to discriminate photoreceptor cell species in xenotransplanted chimeric retinas

**DOI:** 10.1101/2025.03.09.642229

**Authors:** Kang V. Li, Annabelle Pan, Ying V. Liu, Bani Antonio-Aguirre, Joyce Wang, McKaily Adams, Christina McNerney, Sai Bo Bo Tun, Kenneth Jimenez, Yuchen Lu, Zhuolin Li, Minda McNally, Veluchamy A. Barathi, Robert J. Johnston, Mandeep S. Singh

## Abstract

Photoreceptor transplantation is being studied to improve visual function in retinal diseases causing blindness, such as age-related macular degeneration, hereditary eye diseases, and traumatic retinopathy, among others. Preclinical studies often involve the delivery of exogenous human photoreceptor cells into the retinas of animal models. In such experiments, a key readout is the differential frequency of donor cell somatic integration versus artificial labeling secondary to material transfer of cytosolic or nuclear labels from donor to recipient cells. For this analysis, the ability to recognize photoreceptor nuclei as being of donor (human) versus animal is key, but purely immunohistology discrimination can be challenging because of antigenic species overlap or intercellular antigen transfer. To address this challenge, we sought to develop and validate a computational technique to discriminate between photoreceptor cells of different animal species based on machine learning of nuclear morphology. Here, we aimed to evaluate the feasibility of using computer-assisted detection of separate nuclei and employing random forest classification to automate the species differentiation, among DAPI-stained photoreceptors after xeno-transplantation of human photoreceptors into the retinas of mice and pigs.

Our models were trained on single-species samples and validated with mixed-species samples. We then transplanted human embryonic stem cell-derived retinal organoid cells into rodent and pig retinal degeneration models. The random forest model accurately determined cell identity post-xenotransplantation, validated by histological assessment using an anti-human nuclear antibody.

Our results support the potential efficacy of employing machine learning image analysis and classification techniques that may promote experimental rigor, minimize observer bias, and enable high throughput semi-automated workflows for transplantation outcomes analysis. The methodological framework reported here may enable a more nuanced and precise analysis of the behavior of transplanted photoreceptors for the purposes of human retinal regeneration.

## Introduction

Retinal degenerative diseases, including age-related retinal degeneration (AMD), retinitis pigmentosa (RP), and Stargardt disease, among others, are characterized by the degeneration of retinal cells, ultimately leading to irreversible blindness. In 2019, approximately 19.83 million individuals aged 40 years and older in the United States were affected by AMD, while RP and Stargardt disease were reported to have a prevalence rate of one case per every 4,000 and 20,000 people, respectively ^1–3^. Retinal cell transplantation has been envisioned as a potential treatment for treating retinal degenerative diseases.

To pave the way for future clinical trials, preclinical animal transplantation studies are imperative. These studies serve to establish proof-of-concept, develop surgical instruments, optimize surgical procedures, and establish appropriate immunosuppression regimens ^4–7^. However, the behavior of donor cells post-xenotransplantation manifest in potentially unexpected ways. Notably, investigations have demonstrated that in certain experimental condition, fluorescence-labeled donor photoreceptors which appear somatically integrated into the photoreceptor layer of the recipient, are instead recipient photoreceptors that acquired the fluorescent protein by cellular component transfer from the donor cells. This finding has led to a reassessment of the identification and definition of integration within the field ^8–12^. Furthermore, we observed that transplanted retinal organoid-derived cells exhibited migration throughout all retinal layers over considerable distances ^13^. Hence, the precise discrimination between donor and recipient cells is imperative to evaluate cellular dynamics following transplantation. Greater rigor in histological analysis of post-xenotransplantation samples will facilitate the advancement of safe and effective clinical treatment to regenerate retinal photoreceptor cells, with the ultimate aim of improving visual function.

Traditionally, cell identity has been determined based on a limited number of biomarkers, with fluorescent labeling used to differentiate specific cell species within heterogeneous populations. However, this approach may prove inadequate in accurately identifying cells of interest, especially considering the possibility of cellular component transfer (CCT) of labels from donor into recipient cells. CCT may involve donor-derived cytoplasmic, nuclear, and mitochondrial antigens can be transferred to the recipient in retinal cell transplantation ^14^. Furthermore, cross-reactivity involving human-specific nuclear markers was also observed in the transplantation of human-induced pluripotent stem cell (hiPSC)-derived photoreceptors into non-human primates ^15^. In recent years, a rapidly expanding range of high-throughput, single-cell experimental techniques have emerged.

These methods enable a more rigorous definition of cell identity based on a cell’s epigenome, transcriptome, and proteome ^16, 17^. Nonetheless, these technologies require prior knowledge of purified cell populations and access to relatively expensive technology platforms for data acquisition. Alternatively, analysis of cell and nucleus morphology presents a readily accessible method for cell identification. Notably, in retinal cell identification, multiple histologically discernible features have been utilized to distinguish retinal ganglion cells from other retinal cell types ^18^. However, these assessments are typically performed through labor-intensive and time-consuming visual estimation, leading to high inter- and intra-observer variability ^19^. Currently, there is no established automated method for processing and distinguishing photoreceptor species on a large scale in transplantation models.

To address the challenge of identifying the cell species after xenotransplantation, we developed an innovative methodology utilizing computer-assisted detection of cell features that are used in random forest models to classify the post-xenotransplantation species of photoreceptors. Our approach leverages QuPath ^20^, a publicly available semi-automated software for image processing that captures quantitative data on various features of photoreceptor nuclei from diverse sources included C57BL/6J mice, domestic pigs, nonhuman primates, and stem cell-derived retinal organoids. Using these extracted features, we implemented a command-based software tool called RetFM-Class, which integrates the data from QuPath to perform random forest classification of cell types ^18^. In initial performance tests, we applied random forest classification to evaluate the xenotransplantation of human embryonic stem cells derived retinal organoid (hRO) cells into retinal degeneration mice and pigs. Our goal was to evaluate whether this approach could successfully distinguish the identity of hRO photoreceptor nuclei from the recipient photoreceptors of different species, using anti-human nuclear antibody to determine ground truth classification. Through these advancements, we sought to develop a reproducible, rapid, and cost-effective approach to accurately determine the photoreceptors species following xenotransplantation, facilitated by the utilization of machine learning techniques.

## Materials and methods

### Animal

All animal experiments were carried out in accordance with the ARVO Statement for the Use of Animals in Ophthalmic and Vision Research. All procedures were approved by the Johns Hopkins University Animal Care and Use Committee (approval M016M17, SW22M277) and the SingHealth Institutional Animal Care and Use Committee (approval 2015/SHS/1134).

We used *Gnat1^−/−^; Gnat2^−/−^; ROSA^nT-nG^*mice (aged 6 to 8 weeks, n = 6 eyes; *Gnat1^−/−^; Gnat2^−/−^*double-knockout mice were a kind gift from Dr. Marie Burns (University of California, Davis) and Dr. King-Wai Yau (Johns Hopkins University)), domestic pigs (aged 4 to 6 months, n = 6 eyes), and nonhuman primate (aged 6 to 8 years, n =4 eyes) in this study.

### Retinal organoids

Human embryonic stem cells (hESC)-derived retinal organoids are generated from the H9 CRX- tdTomato cell line and CRX-GFP cell line which were generously provided by Dr. David M. Gamm (University of Wisconsin-Madison, USA) and Dr. Donald J. Zack (Johns Hopkins University, USA), with approval from the Johns Hopkins ISCRO (ISCRO00000249). The differentiation process of retinal organoids followed a previously published protocol ^13^. In brief, the hESC were dissociated using Accutase at 37°C for 12 minutes and seeded at a density of 3,000 cells per well in 50 μl of mTeSR1 medium. The cells were seeded into 96-well ultra-low adhesion round bottom Lipidure coated plates (AMSBIO, MA, USA, No.51011610). The cells were initially cultured in hypoxic conditions (10% CO2 and 5% O2) for 24 hours and then moved to normoxic conditions (5% CO2) on day 1. From days 1 to 3, each well received 50 μl of BE6.2 media supplemented with 3 μM Wnt inhibitor (IWR1e, EMD Millipore, MA, USA, No. 681669) and 1% (v/v) Matrigel. From days 4 to 9, 100 μl of media were removed from each well and replaced with fresh media. On days 4-5, BE6.2 media containing 3 μM Wnt inhibitor and 1% Matrigel were added, while on days 6-7, BE6.2 media supplemented with 1% Matrigel were added. On days 8-9, BE6.2 media containing 1% Matrigel and 100 nM Smoothened agonist (SAG, EMD Millipore, No. 566660) were added. On day 10, the aggregates were transferred to 15 mL tubes, washed three times with DMEM (Gibco, No. 11885084), and resuspended in BE6.2 medium with 100 nM SAG in untreated 10 cm polystyrene petri dishes.

Media was subsequently changed every other day. On day 11, retinal vesicles were manually dissected using sharpened tungsten needles. Following dissection, the cells were transferred to 15 mL tubes and washed twice with 5 mL of DMEM. From days 14 to 17, long-term retina (LTR) media supplemented with 100 nM SAG was added. From days 18 to 21, the cells were maintained in LTR media and washed twice with 5 mL of DMEM before being transferred to new plates to remove dead cells. The LTR medium was supplemented with 1 μM all-trans retinoic acid (ATRA; R2625; Sigma) from days 22 to 138. Additionally, 10 μM Gamma secretase inhibitor (DAPT, EMD Millipore, No. 565770) was added to the LTR medium from days 28 to 42. To minimize aggregation, retinal organoids were grown at a low density of 10-20 organoids per 10 cm dish.

### Xenogeneic transplantation

Donor retinal organoid cells, obtained as micro-dissected multilayered retinal fragments, were derived from CRX-tdTomato and CRX-GFP hESC-derived retinal organoids (aged 134 days, n=4). The cultured human retinal organoids were imaged using a fluorescent microscope (Carl Zeiss, Jena, Germany) for reference. The retinal organoids were carefully dissected into micro-dissected fragments using a 27-gauge horizontal curved scissors (VitreQ, Kingston, NH, USA) under a dissection microscope, and then were dissociated into single cells with papain dissociation system (Worthington Biochemical Corporation) for transplantation.

The donor retinal organoid cells were transplanted into the subretinal space of *Gnat1^−/−^; Gnat2^−/−^; ROSA^nT-nG^*mice, following a previously reported protocol ^21^. In brief, recipient mice were anesthetized through intraperitoneal injection of ketamine (100 mg/kg body weight) and xylazine hydrochloride (20 mg/kg body weight). Pupillary dilation was achieved using 1% tropicamide (Bausch & Lomb, Rochester, NY, USA). The mouse corneas were covered with Sodium Hyaluronate (Healon GV, Abbott Medical Optics Inc. CA, USA) and cover glasses (Deckglaser, USA) to facilitate transpupillary visualization. The donor retinal organoid cells were carefully loaded into the 33G microneedle attached micro-syringe (Hamilton, Reno, NV, USA) and tangentially injected into the subretinal space through the sclera of the recipient mice. The success of the injection was confirmed by direct visualization through the dilated pupil of the recipient under a surgical microscope (Leica, Wetzlar, Germany).

Retinal organoid cells were also transplanted into a pig model with hyaluronic acid-induced retinal degeneration. One week prior to transplantation, a subretinal injection of 200 μL of 0.5% hyaluronic acid (Healon) was administered at the nasal superior region of the right eye to induce a mild retinal degeneration. On the day of transplantation, anesthesia was induced using a Ketamine/Xylazine cocktail (20-30 mg/kg ketamine and 2-3 mg/kg xylazine) via intramuscular injection. Propofol (0.83- 1.66 mg/kg) was intravenously administered to facilitate intubation. The depth of anesthesia was assessed based on palpebral and jaw tone. Maintenance anesthesia was achieved using 1-3% isoflurane with mechanical or manual ventilation and 100% oxygen supplementation. Physiological monitoring during surgery included pulse oximetry, end-tidal carbon dioxide, heart rate, and respiratory rate. Topical 1% Tropicamide and 2.5% Phenylephrine were applied to dilate the operative eye, and topical 0.5% Proparacaine Hydrochloride (Tetracaine) was used for local anesthesia. The periocular area was sterilized using a 10% povidone iodine solution, and the surgical field, as well as the eye, were prepared with a sterile drape. Anterior chamber paracentesis was performed to reduce intraocular pressure prior to cell injection. The transplanted cells were loaded into a 1 mL zero dead space syringe and then a 25-gauge needle attached to the syringe tip for achieving the subretinal space. Forceps were used to grip the conjunctival margin adjacent to the limbus to provide counter-traction during needle insertion and maintained the eye at an optimal angle. The needle angle was adjusted to be approximately 15 degrees above the sclera. Slow penetration of the sclera-choroid complex was performed until the needle tip was observed to reach the subretinal space. The retinal cells were then slowly injected into the subretinal space, resulting in the formation of a retinal bleb. The needle was held in place for 1 minute to ensure stable intraocular pressure and minimize reflux before carefully removing it. Intraocular Ozurdex was directly injected into the posterior segment of the eye, and a single dose of Methotrexate (400 µg in 0.1 mL saline), an immunosuppressant, was delivered to the eye. Subconjunctival administration of antibiotics was performed. After 3 weeks, the pigs were euthanized, and enucleation of the eyes was conducted for subsequent histological analyses.

### Histology analysis and Cell Feature Extraction

For WT control and transplantation animals, after euthanasia, enucleated eyes were fixed in 4% paraformaldehyde. Then, eyes were immersed in increasing concentrations of sucrose (10%, 20%, 30%) for dehydration. Cryosections were collected and processed for immunofluorescent labeling with DAPI (4′,6-diamidino-2-phenylindole), and anti-Ku80 antibody (rabbit anti-human, 1:200, ab80592, Abcam). Fluorescence images were acquired with a confocal microscope (Carl Zeiss, Jena, Germany). QuPath was used to automatically obtain four morphological features of the photoreceptor nuclei, namely, the nuclear cross-sectional area, perimeter, circularity, and aspect ratio (AR). AR was determined as the ratio of the nuclei’s major axis to its minor axis.

### Random Forest Classification

The Random Forest (RF) package in R was applied to create multiple binary classifier models according to the variety of species into which ROs were transplanted. Each RF model was trained using QuPath-extracted cell feature data from two species as input and produce a binary species classification as output. RF models are comprised from an ensemble of decision trees, which are each independently trained on a random subset of a fixed proportion of all available training data.

Variability between trees is generated through random feature selection and threshold setting at each node of the trees. The final classification returned by RF is determined through a majority vote among the ensemble of individual decision trees. We used the package “randomForest” in R (version 4.2.1) with default parameters and 500 trees per model. Training sets used information from pre-transplantation individual species’ cells, while test sets consisted of cells after transplant. Ground truth was based upon the presence of human-specific nuclear markers.

### Validation and Evaluation

Out-of-bag (OOB) validation was applied to the models to assess their validity and accuracy. During the construction of each decision tree in the ensemble, a random subset of the training data is used, leaving some observations unused as out-of-bag samples. For each cell in the dataset, the OOB process collects predictions made by trees that did not include this cell in its training set. The majority vote from these out-of-bag trees is considered as the predicted output for that specific observation. By comparing these predictions to the true labels of the cells, the OOB validation efficiently estimates the accuracy of the RF model when generalizing to new, unseen data. In addition to OOB, each model’s performance was assessed using area under the ROC curve (AUC) analysis with test data comprised of each cell type following RO transplantation.

To further characterize our RF models, feature importance analysis was conducted through randomForest::varImpPlot. This method determines the importance of each variable in contributing to classification decisions based on two methods: first, through the reduction in classification accuracy when the variable is left out of permuting predictions, and second, through the total reduction in node impurity (represented with Gini index) when the variable is left out ^22^.

### Statistics and graphs

Statistical analysis was conducted using one-way ANOVA in RStudio (R Core Team, 2020) and Prism Software Version 7 (GraphPad Software Inc., La Jolla, CA). One-way ANOVA was employed to compare the means of each group against the means of all other species. Statistical significance was determined at *p* < 0.05. Bonferroni correction was applied to adjust for multiple comparisons.

Mean differences and 95% confidence intervals were calculated. The cartoons in Figures 1, 4 were created with BioRender.com (agreement number: RG22SB5FYK, QD22SB5ZJ4).

**Figure 1.**
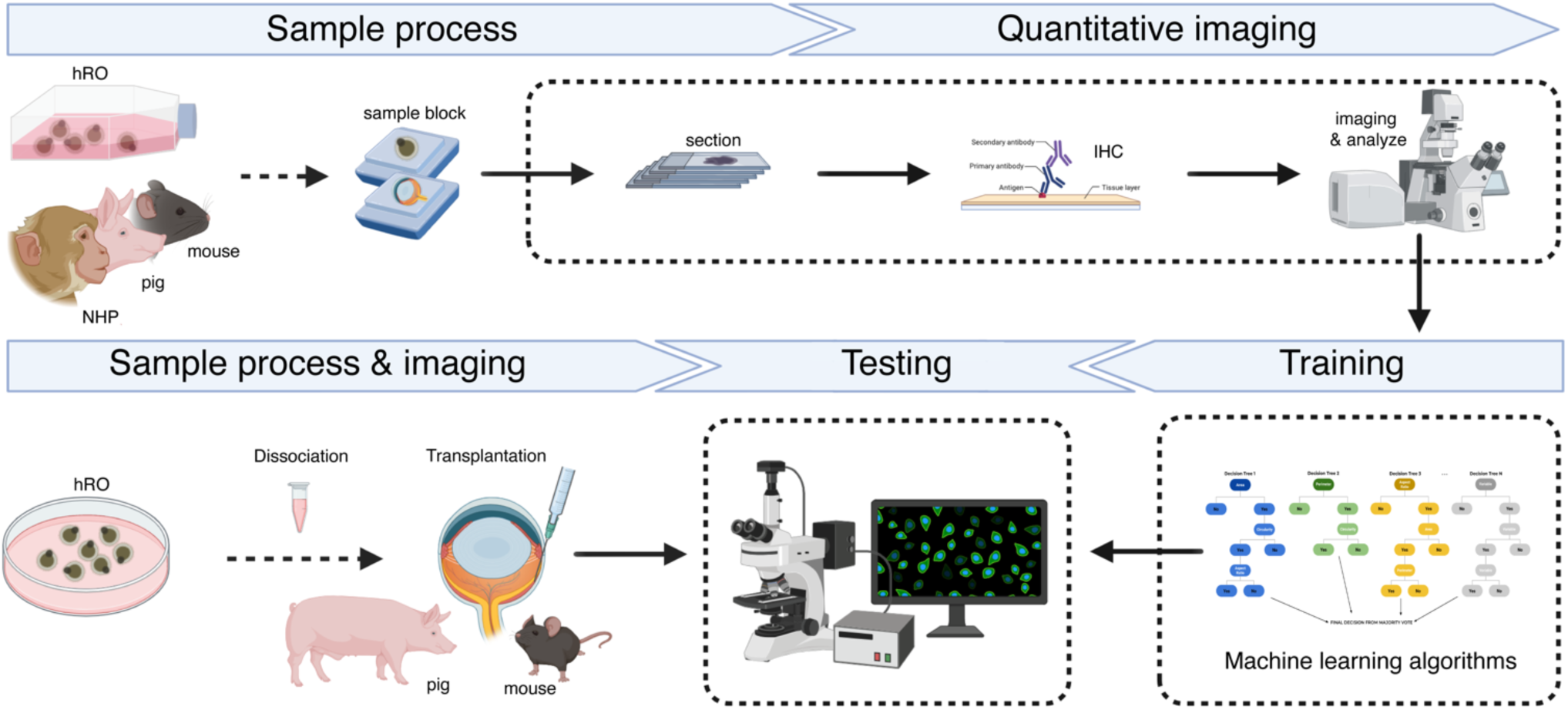
Schematic representation of study methodology. Morphologic data were extracted from histology sections of mouse, pig, nonhuman primate (NHP) retinas, alongside human retinal organoid (hRO) photoreceptor cells. These data were used to train the random forest model. Then, the model was tested using histological slides from two xenotransplant combinations: human-into-mouse and human-into-pig.

## Results

### Photoreceptor nucleus from mouse, pig, nonhuman primate (NHP), and hESC-derived retinal organoids (hRO) present different morphological characteristics

First, we aimed to evaluate morphological differences between photoreceptor cell nuclei from different animals. To do so, we initially gathered data from sections of adult mouse, adult pig, and adult NHP retinas, alongside photoreceptor cells in human retinal organoids (hRO-PRC) (**Figures 1 and 2A**). The photoreceptor layer in the retina sections were assigned using Cone-Rod Homeobox (CRX), which is specifically expressed in developing and mature photoreceptors, and Recoverin (RCVRN), primarily localized in the photoreceptor outer segments of the retina. We employed QuPath software to analyze the morphological characteristics of the photoreceptor nuclei of each different animal. The four morphological characteristics we analyzed were: cross-sectional area, perimeter, circularity, and aspect ratio (AR). We focused on morphological differences that could eventually be used to automatically detect hRO-PRC nuclei (from human donor cells) among surrounding mouse, pig, or NHP photoreceptor nuclei (different transplant recipient animals) recipient animals.

**Figure 2.**
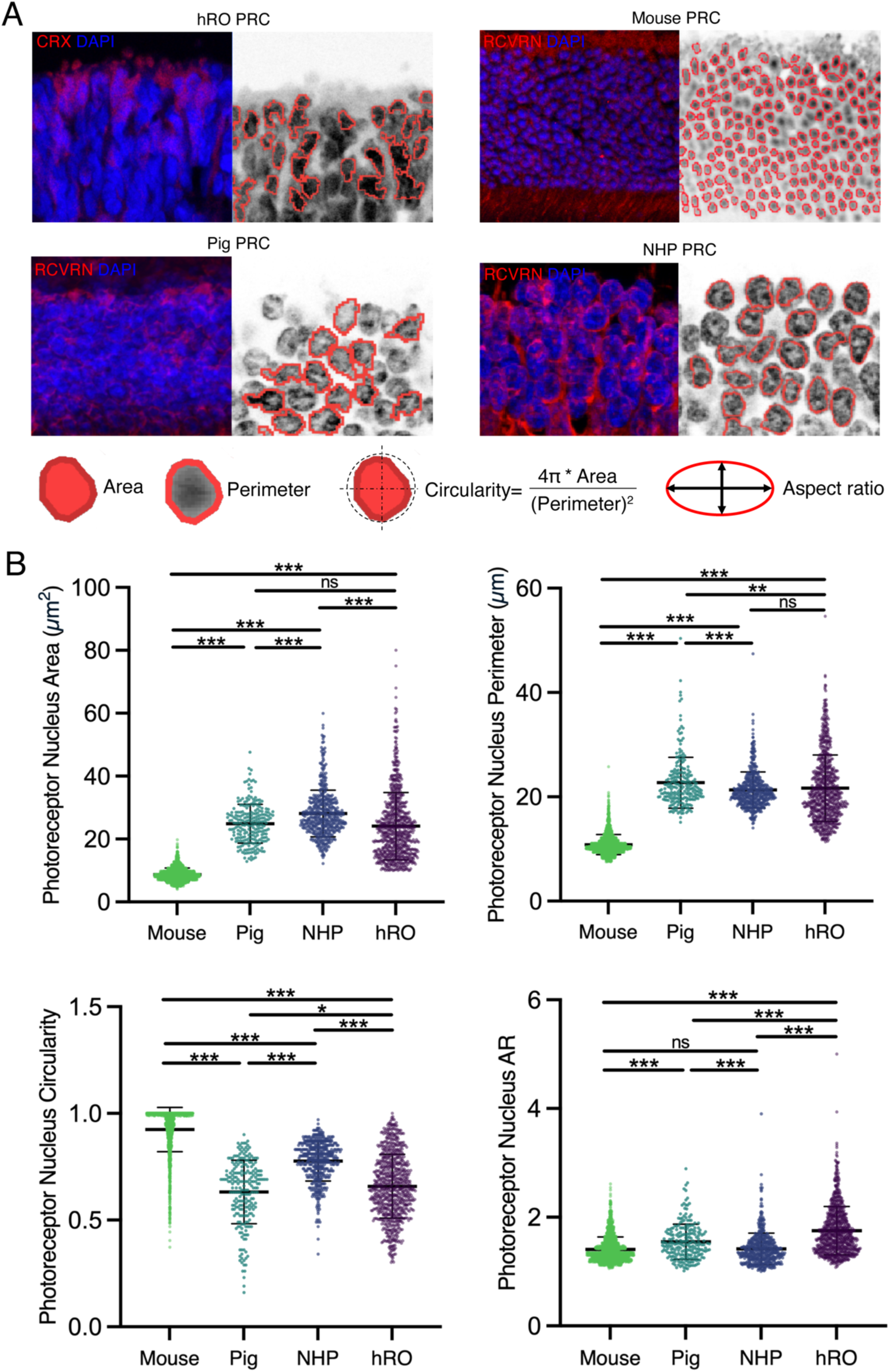
Comparative analysis of mouse, pig, non-human primate (NHP), and human retinal organoid (hRO) photoreceptor nuclear morphology. (A) Cone-Rod Homeobox (CRX) and Recoverin (RCVRN) were used to label photoreceptor cells (PRC), Subsequently, we analyzed nuclear morphology using QuPath software. (B) A total of 1916 mouse PRC, 463 pig PRC, 517 NHP PRC, and 815 hRO PRC were analyzed. We analyzed four nuclear morphology characteristics: two size features (cross-sectional area and perimeter), and two shape features (circularity and aspect ratio [AR]). Overall, we observed several significant differences in these features between mouse, pig, NHP, and hRO PRC. \**p*<0.05, \*\**p*<0.01, \*\*\**p*<0.001

The most common xenotransplant paradigm in this field is to transplant human cells into mouse models, owing to the great diversity of genetic disease models available in this species. With regards to the differentiation of hRO and mouse photoreceptor nuclei (**Figure 2B**), we found that all four morphological characteristics were significantly different. The mean cross-sectional area of human photoreceptor nuclei was significantly greater than that of mouse cells (24.10 ± 10.71 *vs.* 8.68 ± 2.08, *p* < 0.001). Reflecting this size difference, the mean nuclear perimeter of hRO-PRC was also greater than mouse (21.66 ± 6.37 *vs.* 10.84 ± 1.91, *p* < 0.001). In terms of nuclear shape, hRO-PRC showed lower mean circularity than mouse cells (0.66 ± 0.15 *vs.* 0.92 ± 0.10, *p* < 0.001). In accordance with this finding, the mean AR was significantly greater in human than mouse cells (1.75 ± 0.44 *vs.* 1.41 ± 0.23, *p* < 0.001). These data indicate that human embryonic-stage photoreceptors are clearly larger and more ovoid than mouse.

In other xenotransplant experiments, investigators may choose a pig recipient for human donor cell grafts. In this regard, hRO and pig photoreceptor nuclei showed less marked differences in morphological characteristics. In terms of size, the mean cross-sectional area of pig and human photoreceptors were no different. However, the mean nuclear perimeter was significantly greater in pig than human (22.72 ± 4.88 *vs.* 21.66 ± 6.37, *p* = 0.0014), possibly reflecting differences in nuclear membrane boundary infolding pattern. In terms of shape, there was a small but statistically significant difference in mean circularity, with human cells being only slightly more circular (0.63 ± 0.15 *vs.* 0.63 ± 0.15, *p* = 0.0186). The other shape metric, mean AR, was also only slightly (but significantly) greater in human than pig photoreceptor cells.

In a third xenotransplant paradigm, human cells may be transplanted into NHP recipients to address questions relating to, for example, the presence of a recipient macula. In this context it will be important to distinguish between human and NHP photoreceptor cells when studying the histological outcome. Surprisingly, based on the four morphological characteristics we analyzed, hRO and NHP photoreceptor cells showed several key differences. The mean cross-sectional area of hRO photoreceptor cells was significantly less than NHP photoreceptor cells (24.10 ± 10.71 *vs.* 28.12 ± 7.44, *p* < 0.001) but the nuclear perimeter was similar in both species. In terms of shape, human and NHP cells were different. NHP photoreceptors were more circular (mean circularity, 0.78 ± 0.09 *vs.* 0.66 ± 0.15, *p* < 0.001) In accordance with this finding, the mean AR was significantly greater in human than NHP cells (1.75 ± 0.44 *vs.* 1.42 ± 0.29, *p* < 0.001), reflecting a more oval shape of human photoreceptor nuclei. It is worth noting that the highest aspect ratio of the hRO-PRC nucleus could potentially be attributed to their developmental stage around day 120, during which they are still undergoing differentiation into mature photoreceptors.

### Random Forest Model Development

RF models were trained using datasets containing cell features from hRO, mouse, pig, and NHP photoreceptors. For each species we analyzed n=815 hRO photoreceptors, n=1916 mouse photoreceptors, n=463 pig photoreceptors, and n=517 NHP photoreceptors. Results from an out-of-bag validation analysis are shown in Figure 3. The RF model was most accurate in distinguishing hRO-PRC vs. mouse photoreceptor, with an overall accuracy of 93.1%. This was followed by hRO- PRC vs. NHP photoreceptor, 82.4% OOB accuracy, and lastly hRO-PRC *vs.* pig photoreceptor, with 77.9% OOB accuracy (**Figure 3A**). The accuracy was validated by anti-human nuclear antibody. In the model performance analysis, we observed that the sensitivities are comparatively higher than their corresponding specificities for RF models discriminating between hRO-PRC and pig photoreceptor (sensitivity of 82.3% *vs.* specificity of 65.2%) as well as hRO-PRC and NHP photoreceptor (sensitivity of 86.2% *vs.* specificity of 76.6%). Interestingly, a contrasting trend is noted in the discrimination between hRO-PRC and mouse photoreceptor, where the sensitivity (88.4%) is lower than the specificity (95.1%).

**Figure 3.**
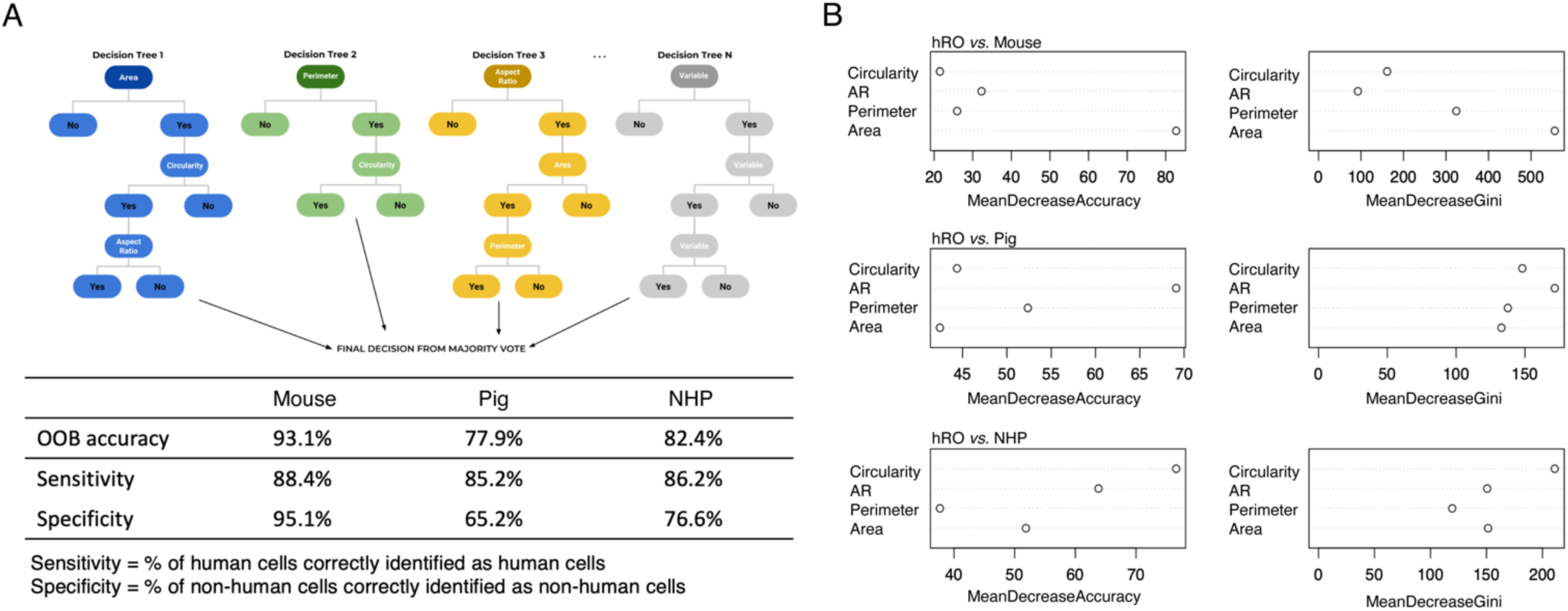
Out-of-bag validation analysis of the trained random forest model. (A) Decision tree and the accuracy, sensitivity, and specificity in discriminating photoreceptors derived from human embryonic stem cell-derived retinal organoids versus those from mouse, pig, and nonhuman primate (NHP). (B) Variable importance plot illustrating the significance of predictors in the random forest model.

As cross-sectional area, perimeter, aspect ratio, and circularity are fundamental factors used to define the shape of a cell nucleus, our models in particular highlighted the importance of cross-sectional area and aspect ratio in accurately classifying between different species of photoreceptor cells. It is noteworthy that the most accurate model, which distinguished between hRO-PRC and mouse photoreceptor, relied most on cross-sectional area (**Figure 3B**), while the models for distinguishing hRO-PRC *vs.* pig photoreceptor and hRO-PRC *vs.* NHP photoreceptor were less accurate and relied most on aspect ratio and circularity. It is possible that the accuracy of the latter two models is limited by the accuracy of calculating an aspect ratio using two-dimensional images that represent a three-dimensional cell, which could have wide-ranging aspect ratios depending on the angle at which it is observed.

### Random Forest Model Testing

We transplanted hRO-PRC into retinal degenerated mouse and pig models and then tested our RF models on cell features obtained from post-transplant cells. NHP transplants were not performed in this study. We analyzed n=11 human cells and n=170 pig cells for hRO cells to pig transplants, n= 9 hRO cells and n=8 mouse cells for human to mouse transplants. The AUC for an RF classifying cells in human to pig transplants was 0.70 (95% CI: 0.58-0.83), and AUC for human to mouse transplants was 1.0 (95%CI: 1.0-1.0) which indicates perfect discriminatory power (**Figure 4A**).

**Figure 4.**
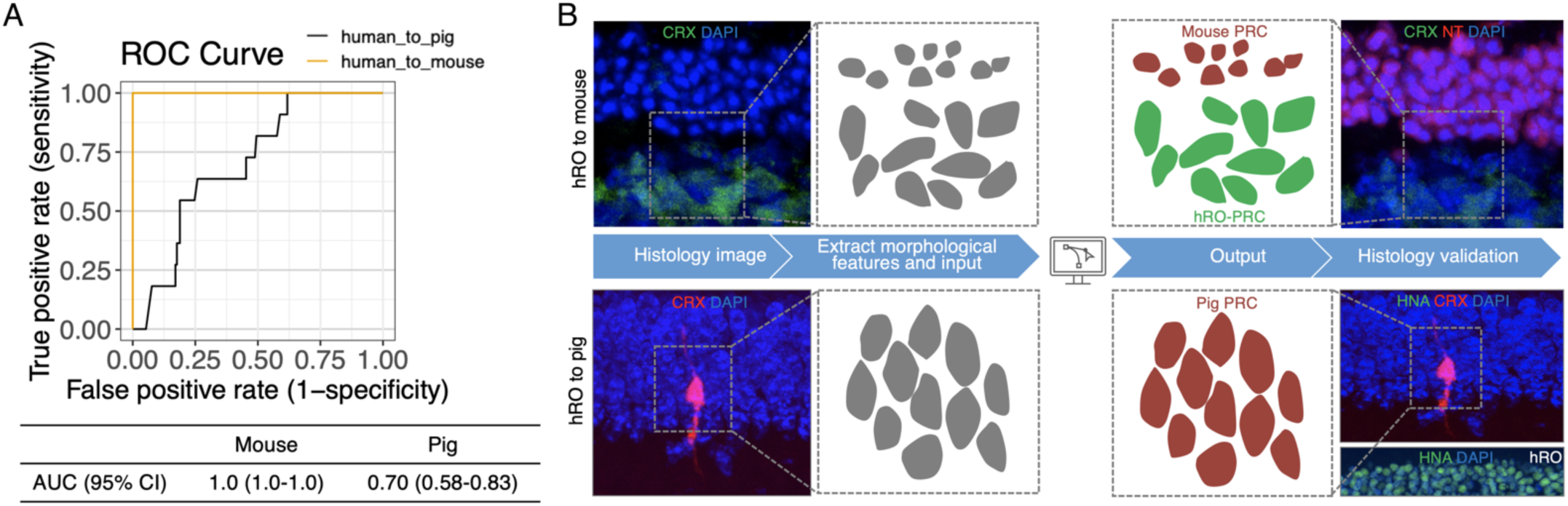
Evaluation of the random forest model in xenotransplantation across animal models. (A) Area under the curve (AUC) for the random forest classifier distinguishing between cells in human-to-pig transplants was 0.70 (95% CI: 0.58-0.83), while AUC for human-to-mouse transplants was 1.0 (95% CI: 1.0-1.0). (B) Examples of photoreceptor species identification include a region containing both donor (human retinal organoid) and recipient (mouse) photoreceptors, and another region where human retinal organoid cells were injected into the subretinal space of a pig with Healon-induced retinal degeneration.

Separately, we transplanted human hRO CRX-GFP photoreceptors into *Gnat1^−/−^; Gnat2^−/−^; ROSA^nT-nG^* mice. A potential region of contact between donor and recipient photoreceptors was identified utilizing our RF model and determined that the hESC-derived CRX-GFP photoreceptors localized beneath the recipient mice photoreceptors (**Figure 4B**). Notably, neither donor photoreceptors migrated nor integrated into the recipient outer nuclear layer (ONL), and no obvious transfer of GFP from donor photoreceptors to recipient photoreceptors was observed in this specific region. In another distinct case, we injected hESC-derived CRX-tdTomato photoreceptors into the subretinal space of a pig with Healon-induced retinal degeneration. Intriguingly, a cell expressing tdTomato was observed within the recipient pig’s ONL. Using the RF model, the tdTomato-positive cell was identified as a pig photoreceptor, suggesting a potential occurrence of cross-species protein transfer from hRO-PRC to pig photoreceptors (**Figure 4B**). The RF model results were subsequently validated through histological evaluation employing an anti-human nuclear antibody. Further evidence is needed to extrapolate the infrequently observed cross-species protein transfer, a phenomenon sporadically noted in both our transplantation results and other studies. Notwithstanding, the RF model demonstrates its capacity to accurately discern cell species in the investigation of post-xenotransplantation neuronal dynamics.

## Discussion

For investigate the donor cell somatic integration and cellular component transfer after transplantation, donor and recipient cell identification is the first and prior thing that need to be done. We developed and validated that the method of machine learning algorithm is capable of overcoming the proposed task-identification of photoreceptors origins from different species after transplantation. The advantages of histology-based random forest classification are that it simultaneously utilizes multiple features to classify cells, requires no special antibodies or probes, yields simultaneous information on multiple species, is rapid and inexpensive, and is readily applicable to high- throughput analyses.

Random forests is a machine-learning technique that offers excellent performance and flexibility in its ability to handle all types of data, for classification of cells ^23^. It works with both categorical and numerical input variables which is less time-consuming on encoding or labeling data which could be convenient to use for our purpose. Random forests have been more and more used in ophthalmology studies (e.g., in diabetic retinopathy classification, predict response of treatment in neovascular AMD, dry eye diagnosis, detect the activity of Graves orbitopathy, and cornea transplant complication evaluation) ^24–28^. To our knowledge, none of retinal cell transplantation study has used random forest in the ophthalmic literature. By leveraging the untapped potential of random forests in the context of retinal cell transplantation, our study adapted this technique to advance research in retinal regeneration.

The ability to differentiate cells originating from distinct species is pivotal in discerning donor cells from recipient cells in transplantation preclinical studies. Precise determination of cell species holds significant importance in the assessment of neuronal survival, maturation, and integration subsequent to the transplantation procedure. Most approaches for cell species identification utilize antibodies, which has potential limitations, especially if the protein that the antibody tag with is capable transferred among cells. One benefit of random forest is its high accuracy thanks to its “wisdom of the crowds”, with respect to the specificity of our algorithm, a great volume of data has been used for both the training and the validation steps, and in all cases the results have been very precise. But one thing we should pay attention is that the appearance of nuclei varies considerably with staining and tissue preparation conditions, as well as with different nuclear types and pathologies. To help encourage this, we have designed our approach to utilize several experienced researchers to ensure the collected images are consistent and high quality to generate the accurate parameters as training dataset. other ways allowing continuing improvement of the approach without recoding software included to enlarge the sample number in the training dataset and add new features as estimators to increase the number of trees so as to improve the accuracy of the model. Another advantage of the histology-based random forest classification method pertains to uniformity. All manual classification methods are prone to higher variation than automated methodologies. Our approach is based on open-source software and easily accessible reagents, making it possible for researchers to analyze the retina and obtain the results in a uniform fashion. Thus, the approach, and its continued evolution, has the potential to be a useful complement to the subjective grading schemes commonly used in photoreceptor transplantation studies.

This study also has limitations. Initially, the study is constrained by the relatively modest sample size in terms of both animal subjects and the number of cells analyzed. Additionally, it is worth noting that there was variation in the absolute values for nuclear size among the retinas subjected to QuPath analysis. It is our working hypothesis that these disparities are likely attributable to the imprecise delineation of auto-segmented nuclear boundaries. Future work will encompass the evaluation of these methodologies across more extensive datasets, incorporating diverse staining techniques.

## Conclusion

We observed distinct morphological variations among photoreceptor nuclei from mouse, pig, NHP, and hRO-PRC. Training a random forest model using these features enabled precise determination of cell species post-xenotransplantation, validated histologically with an anti-human nuclear antibody. Our method offers a consistent, efficient, and cost-effective approach for identifying transplanted photoreceptor species, promising advancements in retinal cell transplantation research.

## Data availability

The data that support the findings of this study are available on request from the corresponding author, MSS. The original contributions presented in the study are included in the article, further inquiries can be directed to the corresponding authors.

## Acknowledgments

This work was funded by the following funding: National Eye Institute (NEI) R01EY033103 (MSS), Foundation Fighting Blindness (MSS), the Shulsky Foundation (MSS), the Joseph Albert Hekimian Fund (MSS), National Eye Institute (NEI) R01EY030872 (RJJ), the BrightFocus Foundation G2019300 (RJJ), the Maryland Technology Development Corporation 2022-MSCRFD-5895 (RJJ), the Juliette RP Vision Foundation (YVL), the Knights Templar Eye Foundation 147042 (YVL), the Wilmer Pooled Professor Fund (PPF) Lutty Grant (YVL), National Eye Institute (NEI) F31EY033187 (CM), Research to Prevent Blindness (unrestricted grant to the Wilmer Eye Institute at Johns Hopkins University), EY001765 (Wilmer P30 Core Grant, Microscopy Module). We thank Dr. David M. Gamm (University of Wisconsin-Madison) and Dr. Donald J. Zack (Johns Hopkins University) for kindly providing CRX-GFP and CRX-tdTomato cell line. We thank Dr. Marie Burns (University of California, Davis) and Dr. King-Wai Yau (Johns Hopkins University) for kindly gift of the *Gnat1^−/−^; Gnat2^−/−^*double-knockout mice. We also thank all the staff members of the Research Animal Resources at the Johns Hopkins University, for their contributions to animal studies.

